# Antigenicity is preserved with fixative solutions used in human gross anatomy: A mice brain immunohistochemistry study

**DOI:** 10.1101/2022.05.06.490908

**Authors:** Ève-Marie Frigon, Mahsa Dadar, Denis Boire, Josefina Maranzano

## Abstract

**Background:** Histology remains the gold standard to assess human brain biology, so *ex vivo* studies using tissue from brain banks are standard practice in neuroscientific research. However, a larger number of specimens could be obtained from gross anatomy laboratories. These specimens are fixed with solutions appropriate for dissections, but whether they also preserve brain tissue antigenicity is unclear. Therefore, we perfused mice brains with solutions used for human body preservation to assess and compare the tissue quality and antigenicity of the main cell populations.

**Methods:** 28 C57BL/6J mice were perfused with: 4% formaldehyde (FAS, N=9), salt-saturated solution (SSS, N=9), and alcohol-solution (AS, N=10). The brains were cut into 40μm sections for antigenicity analysis and were assessed by immunohistochemistry of four antigens: neuronal nuclei (NeuN), glial fibrillary acidic protein (GFAP-astrocytes), ionized calcium binding adaptor molecule1 (Iba1-microglia), and myelin proteolipid protein (PLP). We compared the fixatives according to multiple variables: perfusion quality, ease of manipulation, tissue quality, immunohistochemistry quality, and antigenicity preservation.

**Results:** The perfusion quality was better using FAS and worse using AS. The manipulation was very poor in SSS brains. FAS and AS fixed brains showed higher tissue and immunohistochemistry quality than the SSS brains. All antigens were readily observed in every specimen, regardless of the fixative solution.

**Conclusion:** Solutions designed to preserve specimens for human gross anatomy dissections also preserve tissue antigenicity in different brain cells. This offers opportunities for the use of human brains fixed in gross anatomy laboratories to assess normal or pathological conditions.

**SIGNIFICANCE STATEMENT:** Neuroscientists currently obtain tissue samples from brain banks. Alternatively, a much larger amount of tissue may be obtained from bodies donated to gross anatomy laboratories. However, they are preserved with different fixative solutions that are used for dissection purposes. We ignore if these solutions also preserve antigenicity of the main cell populations, essential in neuroscientific research. This work is the first to show that two solutions currently used in human gross anatomy laboratories preserve sufficient histological quality, in addition to preserve antigenicity of the main cell populations of the mice brain. This work opens the door to the use of human brain tissue obtained from anatomy laboratories in neuroscientific research.

## INTRODUCTION

Histology remains the gold standard method to assess pathological and normal aging changes of the human brain, accurately depicting cellular morphology and the presence of specific antigens (den Bakker, 2017; Durand-Martel, Tremblay, Brodeur, & Paquet, 2010; Filippi et al., 2019; Hoffmann et al., 2011; Pietrzak, Czarnecka, Mikiciuk-Olasik, & Szymanski, 2018). *In vivo* brain samples can only be obtained through biopsies, which provide small amounts of tissue and are very invasive, hence, rarely performed (Quick-Weller et al., 2018). Neuroscientific histology research typically uses *ex vivo* brains obtained from brain banks, as they provide larger tissue samples, but infrequently complete brains (Carlos et al., 2019; Vonsattel, Amaya Mdel, Cortes, Mancevska, & Keller, 2008).

There are presently few active brain banks in Canada (https://www.cbc.ca/news/science/brain-banks-crucial-for-research-clamouring-for-donors-1.757008). As an alternative, brains of bodies donated to human gross anatomy laboratories could expand the availability of tissue to the neuroscientific research community, since there are numerous body donation programs in Canada (https://www.afmc.ca/en/faculties).

Both, brain banks and anatomy laboratories use chemical fixation to prevent the decay of the tissues (Brenner, 2014; McFadden et al., 2019). The two most widely used chemicals for tissue preservation are alcohol and formaldehyde, because of their antiseptic and antibacterial properties, and their capacity to reduce autolysis (Brenner, 2014; Fox, Johnson, Whiting, & Roller, 1985; Musiał, Gryglewski, Kielczewski, Loukas, & Wajda, 2016). However, they increase the cross-linking of proteins affecting the structure of antigens (Birkl et al., 2016; Dawe, Bennett, Schneider, Vasireddi, & Arfanakis, 2009; Eltoum, Fredenburgh, Myers, & Grizzle, 2001; Im, Mareninov, Diaz, & Yong, 2019; Puchtler & Meloan, 1985; van Essen, Verdaasdonk, Elshof, de Weger, & van Diest, 2010). Additionally, formaldehyde distorts (Mouritzen Dam, 1979; Weisbecker, 2011) and hardens the tissue, making it less flexible (Hayashi et al., 2014; Richins, Roberts, & Zeilmann, 1963; Tolhurst & Hart, 1990), and it is a health hazard (Fisher, 1905; Musiał et al., 2016; Raja & Sultana, 2012). Therefore, high concentrations, such as that used in brain banks and animal fixation (i.e. 4%) (McFadden et al., 2019; Yamaguchi & Shen, 2013) are avoided in human anatomy (Brenner, 2014).

Consequently, human anatomists have developed alternative fixative solutions that combine various chemicals, allowing lower concentrations of formaldehyde. For example, a salt-saturated solution preserves the bodies’ flexibility, optimizing dissection procedures (Coleman & Kogan, 1998; Hayashi et al., 2014). Another example is a solution of multiple alcohols tailored to perform neurosurgical simulations, since the density and retraction properties of the tissue remain closer to the *in vivo* brain (Benet, Rincon-Torroella, Lawton, & González Sánchez, 2014). Despite the good tissue preservation and dissection properties provided by salt and alcohol solutions, it is unclear whether they preserve cellular morphology and antigenicity of the main brain cell populations (Benet et al., 2014; Lyck, Dalmau, Chemnitz, Finsen, & Schrøder, 2008).

Finally, in gross anatomy laboratories, the fixation/embalming procedures are performed by arterial perfusion within 48 hours following death (Brenner, 2014; Frølich, Andersen, Knutsen, & Flood, 1984; Maranzano et al., 2020; Nicholson, Samalia, Gould, Hurst, & Woodroffe, 2005). This inconstant post-mortem delay introduces a variable degree of clotting, which combined with vascular stenosis may negatively impact the perfusion, confounding the fixation quality (Turkoglu et al., 2014).

In this study, we avoided the post-mortem delay by using mice brains, which are fixed immediately by perfusion of the deeply anesthetized animals. We then assessed the overall histology procedures and quality (i.e. perfusion quality, ease of manipulation of the slices and tissue quality), as well as antigenicity preservation (assessed by immunohistochemistry of four antigens) using three solutions. Since 4% formaldehyde (FAS) is the fixative of choice of brain banks and animal fixation protocols, we hypothesized that the histology quality and antigenicity would be superior in mice brains fixed with FAS than those fixed with a salt-saturated solution (SSS) or an alcohol solution (AS) used in human gross anatomy (Benet et al., 2014; Coleman & Kogan, 1998; McFadden et al., 2019; van Essen et al., 2010).

## METHODS

### Experimental design and statistical analysis

We designed an experimental randomized and non-blinded study to compare the variables across three groups. Our control group included the FAS-fixed specimens, and the two experimental groups included the SSS and AS-fixed brains (details in Population section). The categorical variables were compared using Chi-square tests (details in Variables of interest section). All the statistical analyses were corrected for multiple comparisons using Bonferroni correction and were performed using SPSS statistics software (28.0.0 version).

### Population

We used 28 six months old C57BL/6J mice (14 females, 14 males) (Table 1) raised in an enriched environment with controlled ventilation (45 to 60% humidity) and temperature (20 to 25°C) on a twelve-hour daylight schedule. We followed the guidelines of the Canadian Council on Animal Care, and all procedures were approved by the Ethics Committee of the Université du Québec à Trois-Rivières.

**Table 1.**
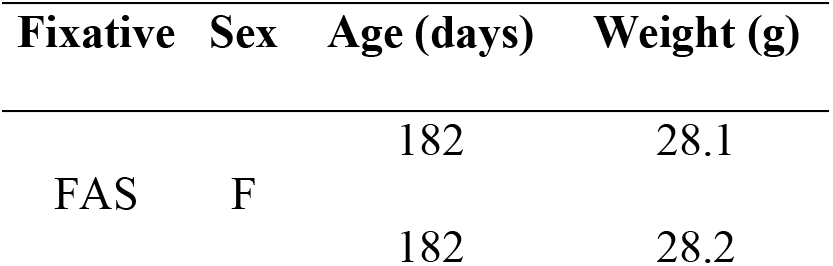
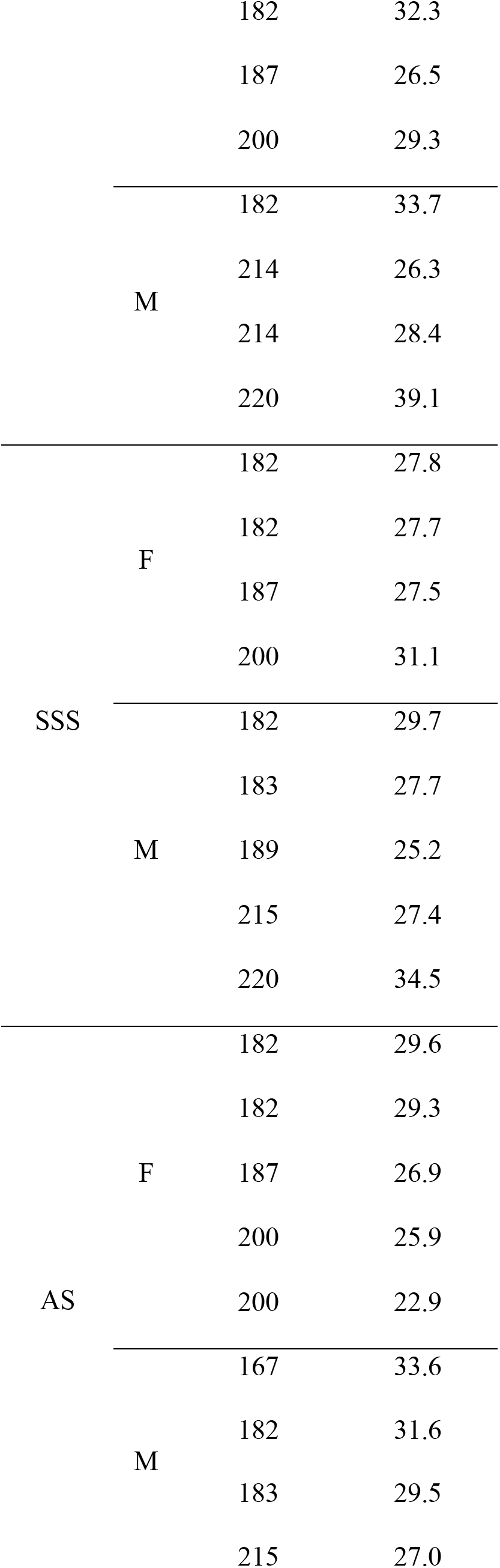
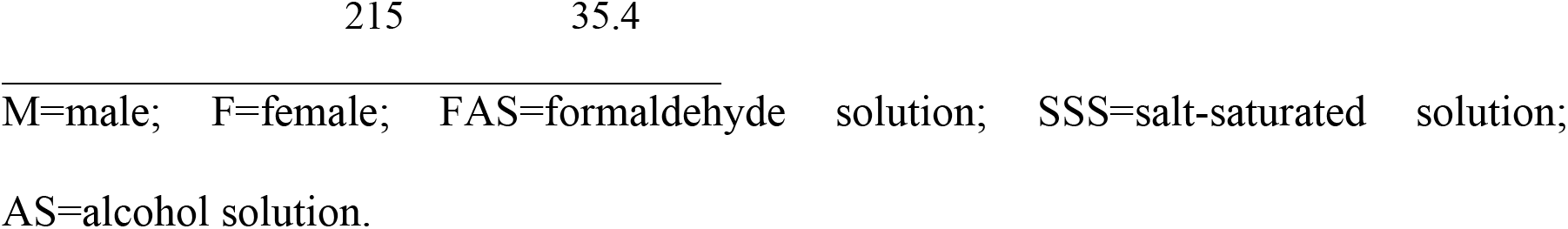
Mice data

### Fixation procedure

Mice were sacrificed by transcardiac perfusion with phosphate-buffered saline (PBS) (0.1M; 0.9% NaCl), followed by a five-minute injection of one of the following solutions (randomly assigned): 1) FAS (N=9), 2) SSS (N=9), and 3) AS (N=10) (Table 2). The perfusion was performed using a gravity technique (Au-Rana, Au-Massa, & Au-Chen, 2022; Paul, Beltz, & Berger-Sweeney, 2008), with the same caliber tubes (15 drops/ml) and cannulas. FAS and SSS bags were placed at the same level, obtaining a similar continuous fluid flow. Due to the higher viscosity and density of the AS, owing to its greater concentration of glycerol, the AS bag was placed 50 centimeters higher to obtain a similar continuous flow than that of the other solutions. We then extracted and post-fixed each brain in the same solution for one to two hours (according to the brain color, which indicates the amount of remaining blood: 2 hours for pink brains, 1.5 h for heterogeneous color brains, 1 h for beige brains). We then immersed the brains in 30% sucrose in 0.1M PBS for cryoprotection overnight. Finally, the specimens were frozen in dry ice and stored at −80°C until use.

**Table 2.** Fixative’s components

| FAS | SSS (Coleman & Kogan, 1998) | AS (Benet et al., 2014) |
| --- | --- | --- |
| 4% formaldehyde | 0.296% formaldehyde | 0.851% formaldehyde |
| PBS 0.1M | 36% NaCl | 62.4% ethanol |
|  | 0.72% phenol | 10.2% phenol |
|  | 2% glycerol | 17% glycerol |
|  | 16% isopropyl alcohol |  |

### Histology processing

Brains were cut using a cryostat (Leica CM1950) in 40 μm coronal sections at −19°C and rinsed three times for five minutes in 0.1M PBS. We then incubated the floating sections thirty minutes in an aqueous solution of 20% methanol and 0.5% H2O2 to quench endogenous peroxidase. Sections were rinsed again three times for five minutes in 0.1M PBS before a two-hour incubation at room temperature in blocking solution (3% Normal Donkey Serum; 0.5% Bovine Serum Albumin; 0.3% Triton X-100 in 0.1M PBS). Then, we incubated the sections overnight at 4°C in the same blocking solution with primary antibodies (Table 3). We rinsed again three times for five minutes in 0.1M PBS before incubation of the sections for two hours in donkey anti-rabbit biotinylated secondary antibody (1:500) at room temperature in the same blocking solution. Sections were rinsed again (3×5minutes in 0.1M PBS) and incubated in an avidin-biotin complex (ABC) kit for thirty minutes in the dark (Vector Laboratories, catalog#VECTPK6100, see fabricant for utilisation procedure). We rinsed three times for five minutes in TRIS-buffered saline (TBS) (0.05M; 0.9% NaCl) before the final incubation in 0.07% diaminobenzidine (DAB) (Sigma, catalog #D5905-50TAB) with 0.024% H_2_O_2_ in TBS for ten minutes. We rinsed the slices in 0.01M phosphate-buffered (PB) before mounting on 2% gelatinated-slides. Finally, sections were dehydrated in alcohol baths (5 minutes each: 70% ethanol, 95% ethanol, 2X 100% ethanol and 2X xylene) and coverslipped using Eukitt.

**Table 3.** Antibodies

| Primary antibodies | Animal | Laboratory | Dilution | Code |
| --- | --- | --- | --- | --- |
| Anti - Neuronal nuclei (NeuN) | Rabbit |  | 1 :3000 | Ab177487 |
| Anti - Glial fibrillary acidic protein (GFAP) | Rabbit |  | 1 :1000 | Ab68428 |
| Anti - Myelin proteolipid protein (PLP) | Rabbit | Abcam | 1 :3000 | Ab254363 |
| Anti - Ionized calcium binding adaptor molecule1 (Iba1) | Rabbit |  | 1 :1000 | Ab178847 |
| Secondary antibody | Animal | Laboratory | Dilution | Code |
| Anti-Rabbit Biotinylated | Donkey | NovusBio | 1 :500 | NBP1-75274 |

### Variables of interest

#### 1) Perfusion quality

we used 3 criteria to assess this variable:

##### 1.1) Signs of perfusion through the right ventricle

this will result in a poorer perfusion since the fixative goes into the pulmonary circulation and indirectly to the brain instead of accessing the systemic (left) circulation, which goes directly to the brain. This was assessed since the higher the density of a solution (namely the AS with high concentration of glycerol), the greater the potential for damage of the thin interventricular septum, with the unintentional passage of the solution from the left to the right ventricle. The indicative signs of perfusion in the right ventricle, used for the score, were red liver, white lungs, and fluid flowing through the nose.

Scores: 3 signs=0, 2 signs=1, 1 sign=2, 0 sign=3.

##### 1.2) Brain color

it is an indication of the quality of perfusion, where the pink color indicates remaining blood in the brain, showing that the perfusion was less effective._In these cases, brains were post-fixed for a longer period to better our odds of a good histology.

Scores: Pink=0 (Figure 1N), Heterogenous=1 (Figure 1M), Beige=2 (Figure 1L).

**Figure 1.**
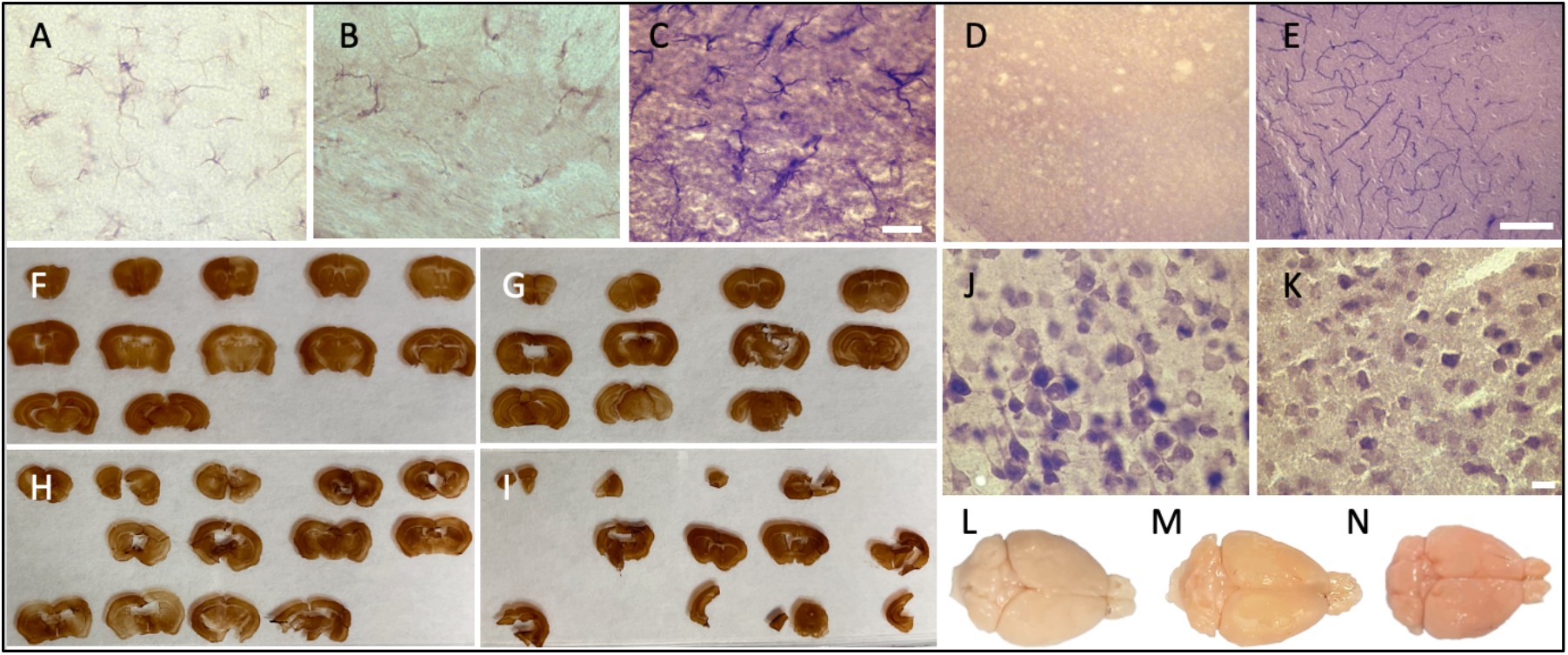
Photomicrographs of the different categorical variables that were assessed. Photomicrographs (60X) of the background intensity (A) light=2; (B) intermediate=1; (C) dark=0, scale bar=20μm. D) Photomicrograph of the presence of dilated vessels (20X) (perfusion quality) and E) of peroxidase vessels (20X) (histology quality), scale bar=100μm. Ease of manipulation (F) very good=3; (G) good=2; (H) poor=1; (I) very poor=0. Photomicrographs (60X) of a uniform neuropil and regular cell contours (J), and a fissured and irregular neuropil and cell contours (K) scale bar=10μm. Brain color (perfusion quality) (L) beige=2; (M) heterogenous=1; (N) pink=0.

##### 1.3) Dilated vessels

a higher injection pressure during the perfusion made the vessels burst, negatively impacting the subsequent microscopic analysis, since the antigens may be masked out by these dilated vessels. This was assessed in the NeuN sections of the specimens.

Scores: Presence=0 (Figure1D), Absence=1.

#### 2) Ease of manipulation

we used 4 signs to assess this variable, namely 1-slice consistency (soft or firm), 2-presence of tears, 3-slices sticking to the brush that was used to manipulate/mount the slice, and 4-presence of rolling slices.

Scores: Very poor (4 signs present=0) (Figure 1I), Poor (3 signs present=1) (Figure 1H), Good (1 or 2 signs present=2) (Figure 1G) or Very good (No sign present=3) (Figure 1F)

#### 3) Tissue quality

we used 3 criteria to assess this variable

##### 3.1) Neuropil quality

a fissured neuropil reflects degradation of the tissue. This was assessed in the NeuN sections of the specimens.

Scores: Fissured=0 (Figure 1K), Uniform=1 (Figure 1J).

##### 3.2) Cell shape

shrunken and shrivelled contours (irregular) are signs of dehydration of the neurons. This was assessed in the NeuN sections of the specimens.

Scores: Irregular (shrunken/shrivelled)=0 (Figure 1K), Regular (smooth)=1 (Figure 1J).

##### 3.3) Peroxidase labelled vessels

the presence of endogenous peroxidase in the endothelial cells of the vessels shows that the quenching procedure was not sufficient for the specimen. This criterion was assessed because it may negatively impact the subsequent microscopic analysis by masking out the antigens. This was assessed in the GFAP sections of the specimens.

Scores: Presence=0 (Figure 1E), Absence=1.

#### 4) Antigenicity preservation

no labeling would show the destruction and/or too much cross-linking of this specific antigen by the fixative. This was assessed for every antigen.

Scores: Absence=0; Presence=1.

#### 5) Immunohistochemistry quality

we used 2 criteria to assess this variable:

##### 5.1) Antibody penetration

it shows the permeability of the tissue. The incomplete condition means the middle of the slice was not labelled, and the complete depicts full penetration at all depth levels. This was assessed for every antigen.

Scores: Incomplete=0, Complete=1.

##### 5.2) Tissue background

the background reflects the quality of the chemical reaction of the DAB labeling. This was assessed for every antigen.

Scores: Dark=0 (Figure 1C), Intermediate=1 (Figure 1B), Light=2 (Figure 1A).

### Microscopy

We assessed variables 3, 4, and 5 using a brightfield microscope (Olympus BX51W1) controlled by Neurolucida software (MBF Bioscience). EMF graded these variables by looking through the full thickness of sections immunolabeled for the four antigens with 60X objective (60X, UPlanSApo 60x/1.40 Oil∞/0.17/FN26,5 UIS2).

## RESULTS

We started by assessing the perfusion quality, finding a significantly higher number of cases without signs of perfusion through the right ventricle in the FAS-fixed specimens (p=0.0002) (Figure 2A). The SSS and AS show a higher number of cases with one sign and three signs of perfusion in the right ventricle respectively, but this observation did not retain significance after Bonferroni correction. We did not find significant differences in brain color, nor dilated vessels across the three fixatives (Figure 2B and 2C). However, pink colored brains were only observed when fixed with AS (N=3) and SSS (N=2). Also, although not statistically significant, there were fewer AS specimens showing dilated vessels.

**Figure 2.**
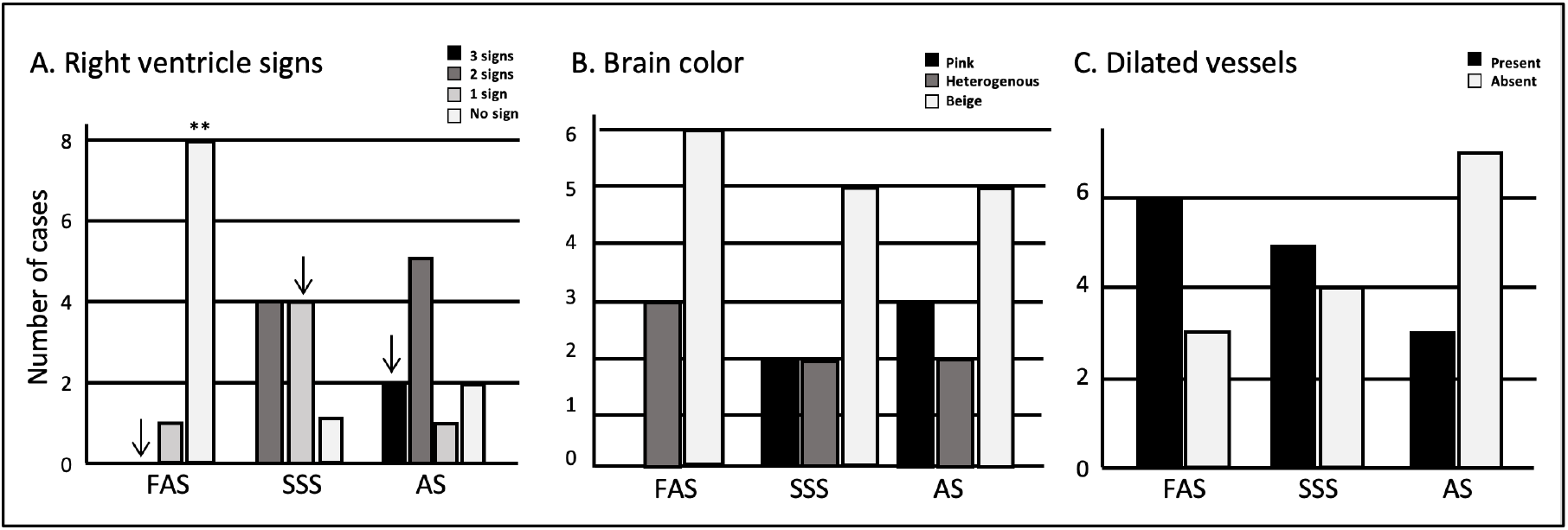
Bar Charts of the perfusion quality. **A)** Signs of perfusion in the right ventricle; **B)** Brain color; **C)** Presence or absence of dilated vessels. Significance after Bonferroni correction: **p<0.001; ↓=significant before Bonferroni correction; FAS=formaldehyde solution; SSS= salt-saturated solution; AS=alcohol solution.

We then assessed the ease of manipulation, finding that the scores obtained for the brains fixed with SSS were significantly lower (p=0.002) than those of the brains treated with the other two solutions (Figure 3). We also found that the scores of the manipulations corresponded more often to a poor manipulation of SSS slices, to a very good manipulation of FAS slices, while the AS had good scores, but these observations did not retain significance after Bonferroni correction.

**Figure 3.**
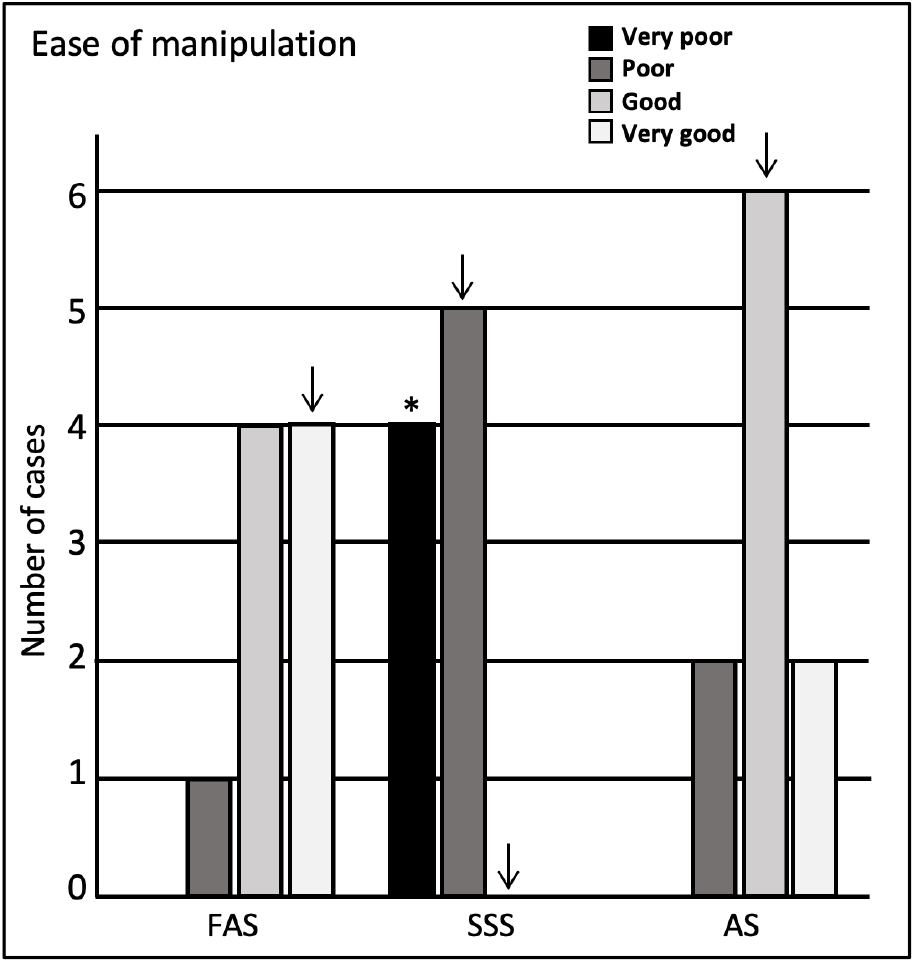
Bar Charts of the ease of manipulation. Very poor=4 signs; poor=3 signs; good=1 or 2 signs; very good= no sign. Significance after Bonferroni correction: *p<0.05; ↓=significant before Bonferroni correction; FAS=formaldehyde solution; SSS=salt-saturated solution; AS=alcohol solution.

Regarding the tissue quality, we found that the neuropil was significantly more often uniform in FAS-fixed specimens (p<0.001). It was significantly more often fissured in the SSS-fixed brains (p=0.007); AS-fixed brains also showed more specimens with a fissured neuropil, which did not retain significance after Bonferroni correction (Figure 4A). Regarding the cellular shape, the specimens fixed with SSS showed significantly more often irregular cells (p=0.004) compared to the brains fixed with AS or FAS (Figure 4B). The FAS-fixed specimens showed mostly regular cells, but this did not retain significance after Bonferroni correction. Finally, we found no significant differences for the occurrence of peroxidase labeled vessels across the three fixatives (Figure 4C).

**Figure 4.**
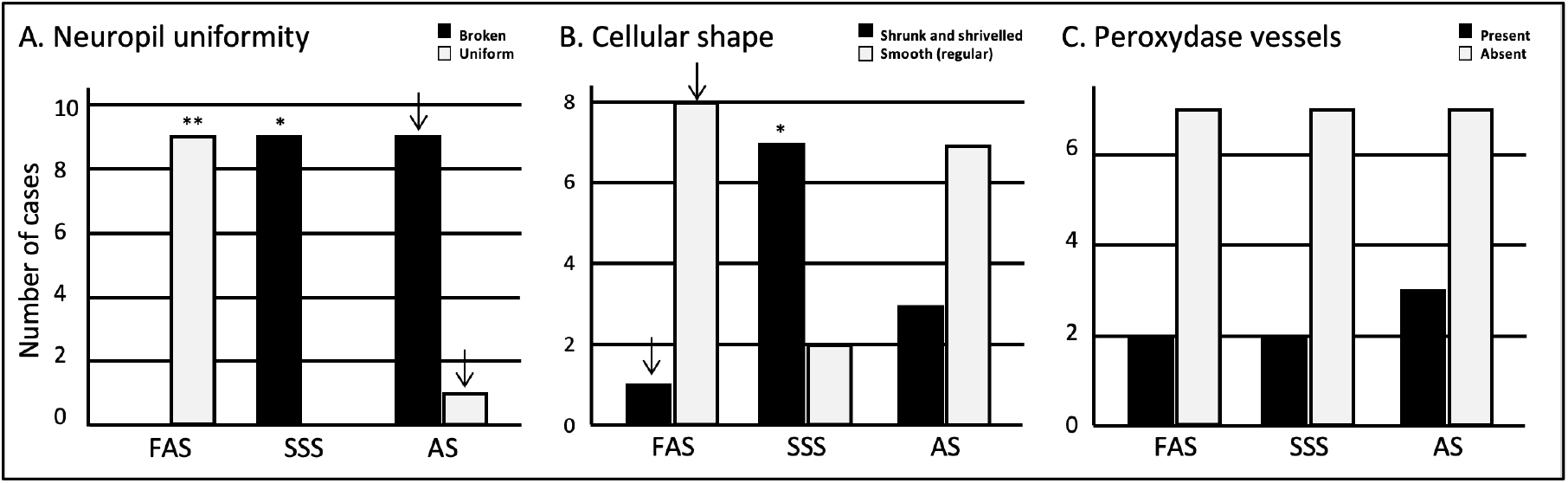
Bar Charts of the tissue quality. **A)** Neuropil uniformity; **B)** Cellular shape; **C)** Presence or absence of peroxidase vessels. Significance after Bonferroni correction, *p<0.05, **p<0.001; ↓=significant before Bonferroni correction; FAS=formaldehyde solution; SSS= salt-saturated solution; AS=alcohol solution.

We also found that the three fixatives preserved antigenicity for the four antigens of interest. Neurons (NeuN) were labeled and showed the neuronal cell bodies (Figure 5A, 5E and 5I). Astrocytes (GFAP) (Figure 5B, 5F and 5J) and microglia cells (Iba1) (Figure 5C, 5G and 5K) clearly showed cell bodies and fine cellular processes. Finally, PLP clearly labeled myelinated fibers (Figure 5D, 5H and 5L) in all the specimens. The four tested antigens were present and comparable in every mouse and therefore, there were no differences between the three fixatives.

**Figure 5.**
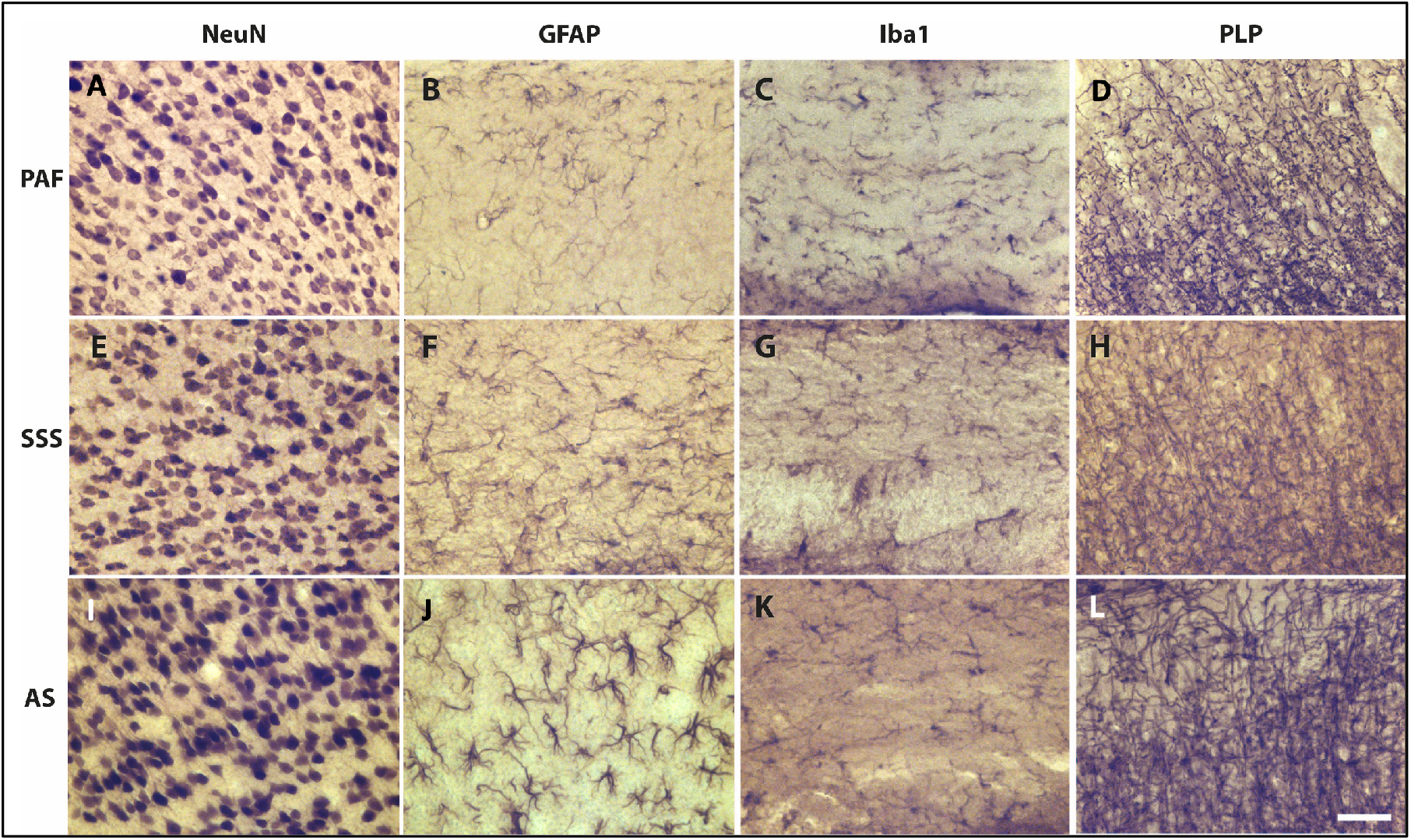
Photomicrographs (40X) of the different antigens of brain samples fixed with three different solutions. **A-E-I)** Neuronal bodies (NeuN), **B-F-J)** Astrocytes (GFAP), **C-G-K)** Microglia (Iba1) and **D-H-L)** Myelin fibers (PLP) fixed with FAS, SSS and AS, respectively. Scale bar = 100μm.

Since all the antigens were present, we were able to assess the quality of immunolabeling using two criteria. First, we assessed the antibody penetration in the depth of the sections for the four antigens. NeuN penetrated completely in all the specimens fixed with AS, while FAS-fixed brains showed three specimens and SSS one specimen with incomplete penetration (Figure 6A). GFAP penetrated completely in the FAS and AS-fixed brains, while three SSS specimens showed incomplete penetration (Figure 6B). These results were statistically significant only before Bonferroni correction. We did not find any differences in the penetration of Iba1 nor PLP between the three fixatives (Figure 6C). However, PLP showed the poorest antibody penetration, especially in FAS-fixed brains (7 incomplete penetration), with AS-fixed brains showing the best PLP antibody penetration (5 complete penetrations) although this was not statistically different across the three solutions (Figure 6D).

**Figure 6.**
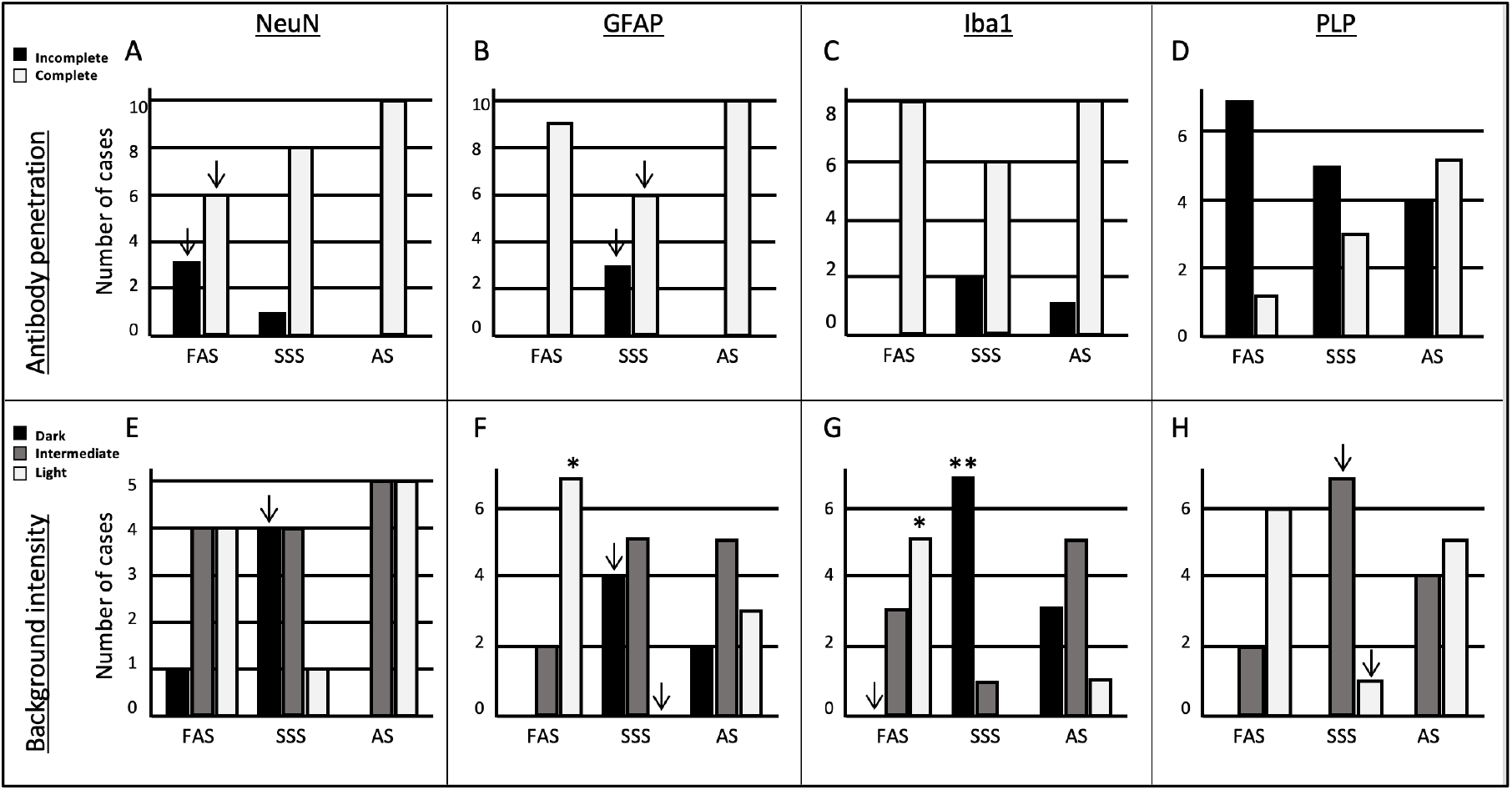
Bar Charts of the immunolabeling quality. Antibody penetration of **A)** NeuN, **B)** GFAP, **C)** Iba1, **D)** PLP. Tissue background of **E)** NeuN, **F)** GFAP, **G)** Iba1 and **H)** PLP. Significance after Bonferroni correction: *p<0.05, **p<0.001; ↓ =significant before Bonferroni correction; FAS=formaldehyde solution; SSS= salt-saturated solution; AS=alcohol solution.

Finally, we assessed the background intensity in slices labeled with the four antigens. The NeuN background labeling was more often darker in brains fixed with SSS, but it did not retain significance after Bonferroni correction (Figure 6E). The GFAP background was significantly more often lighter in brains fixed with FAS than the two other solutions (p=0.001). Additionally, SSS-fixed brains showed a higher occurrence of dark backgrounds and absence of light backgrounds of GFAP than the two other fixatives, but not retaining significance after Bonferroni correction (Figure 6F). In Iba1 labeled slices, we found significantly more often dark backgrounds in the specimens fixed with SSS (p=0.0009), while more light backgrounds were present in FAS-fixed brains (p=0.002). We also never observed dark backgrounds in the specimens fixed with FAS, but this did not retain significance after Bonferroni correction (Figure 6G). In the PLP labeled slices, we found no dark backgrounds for any of the specimens. The SSS-fixed brains present more often an intermediate background and less often a light background than the two other solutions, but this did not retain significance after Bonferroni correction (Figure 6H).

## DISCUSSION

In this study, we investigated the impact of three different fixative solutions, one routinely used in brain banks and animal fixation studies, as well as two solutions used in human gross anatomy laboratories, on the histological quality of brain tissue immunostained for common antigens used in neuroscience research. Our goal was to determine whether the fixatives used in human anatomy procedures could produce histological sections of sufficient quality and preserve antigenicity when administered by perfusion without the confounding effect of post-mortem delay.

We first assessed the perfusion quality since it is the process by which the fixative is delivered to the organ, and potential differences in the flow through certain vessels could compromise the quality of the fixation. Specifically, we foresaw that the AS might be more difficult to perfuse because it contains a high concentration of glycerol (17%) (Benet et al., 2014), which has a higher density and viscosity (Ferreira et al., 2017; Takamura, Fischer, & Morrow, 2012). This caused the thin interventricular septum of the mouse to be broken more often, decreasing the perfusion quality of the AS specimens, which showed a higher frequency of signs of perfusion in the right ventricle, although not statistically significant. As expected, brains were easier to perfuse with FAS, showing significantly fewer signs of right ventricle injection (Winkelman & Beenackers, 2000).

Also, as anticipated, the AS-fixed brains were more often pink, reflecting a higher amount of blood in the brain after perfusion, hence a jeopardized perfusion (note that we did not exclude any of the pink-colored specimens; instead, we treated them with longer postfixation times in the corresponding fixative solution and proceeded to assess the histology variables). Finally, we observed that AS-fixed brains were less prone to showing dilated vessels, which was un unexpected finding considering that the perfusion was done using a higher bag placement of a higher density solution, hence a higher pressure. We speculate that not the density, but the higher viscosity, which determines a slower and weaker flow through the brain vessels, is responsible for the less frequent occurrence of dilated vessels (Ferreira et al., 2017; Takamura et al., 2012; Wilkes, 2017). Alternatively, since dilated vessels may arise as a freezing artifact, the less frequent occurrence in AS specimens could be related to a protective effect against this freezing artifact determined by alcohol, namely glycerol (Bhattacharya, 2018; Kar, Chourasiya, Maheshwari, & Tekade, 2019; Zhang et al., 2020).

We then assessed the ease of manipulation, which determines the ability to cut and mount a full slice, required for an accurate histology analysis. Softer tissue is more fragile, hence very difficult to cut into a full slice with a cryostat. Also, viscous tissue sticks to the brush used to manipulate the slices, which enhances tears or loss of tissue. In this regard, we found that the brains fixed with SSS were the most difficult to manipulate and the FAS-fixed slices were the easiest. This is in agreement with studies on human brains showing that FAS increases the rigidity of the tissue, while SSS-fixed brains are softer (Balta, Cronin, Cryan, & O’Mahony, 2015; Brenner, 2014; Coleman & Kogan, 1998; Eltoum et al., 2001; Fox et al., 1985; Haizuka et al., 2018; Hayashi et al., 2014; Kiernan, 2000; McFadden et al., 2019; Musiał et al., 2016; Nicholson et al., 2005; Weisbecker, 2011). Fortunately, even in the brains that were very soft, and most difficult to manipulate, we could still obtain analyzable slices for all the mice. Moreover, we found that the manipulation of AS-fixed brains was mostly good, and this means the fixative is sufficiently effective to produce tissue with a texture suitable for histology manipulation despite the poorer perfusion results that we obtained.

Regarding the tissue quality, we assessed the neuropil and cellular shape since these characteristics reflect the changes of cell morphology that may be impacted by the different chemicals used for fixation (Spencer, 2017; Troiano, Ciovacco, & Kacena, 2009). We speculate that the shrunken aspect of the neuronal cells bodies observed in SSS brains, but not in the FAS nor AS-fixed specimens, is related to the high concentration of isopropylic alcohol, which is a potent dehydrating fluid, only present in the SSS (Viktorov & Proshin, 2003). Additionally, isopropylic alcohol could have also affected the higher occurrence of fissured neuropil in these brains. In the AS-fixed brains, the higher frequency of fissured neuropil may be due to the high concentration of ethanol, also used as a dehydrating agent (Viktorov & Proshin, 2003). However, despite the poorer tissue quality of AS and SSS-fixed brains, all specimens clearly showed the targeted cells (Figure 5).

In addition to morphological tissue characteristics, we were mostly interested in assessing the presence or absence of antigens in the brains. The detection and quantification of antigens present in the main cells of the brain are essential in a wide range of topics of neuroscientific research, from demyelinating diseases to neurodegenerative processes (Durand-Martel et al., 2010; Filippi et al., 2019; Hoffmann et al., 2011; Pietrzak et al., 2018). Our results showed that antigenicity for all the targets: neuronal nuclei (neurons), glial fibrillary acidic protein (astrocytes), ionized calcium binding adaptor molecule1 (microglia), and myelin proteolipid protein (oligodendrocytic myelin), was equally preserved in brains fixed with any solution (Figure 5), opening the door to the use of brains fixed with solutions currently reserved for gross anatomy purposes.

Finally, we assessed two aspects of the immunolabeling quality that may impact the antigen detection. First, the background labeling, since the darker this background, the lower the contrast with the cells of interest that we aim to visualize, potentially, under-detecting them. Our work showed that SSS brains had the darkest backgrounds, but this did not preclude, in any specimens, the identification of the targeted antigens. Second, the penetration of the antibodies in the full thickness of the slice, which will determine the visualization of the cells at any depth level. This penetration represents the permeabilization of the tissues, differently modified according to the chemical treatment. We found that the AS-fixed brains consistently showed better penetration for all antigens, including the suboptimal PLP, which was relatively less suboptimal with AS than with SSS and FAS. We speculate that this could be related to the chemical mixture of the AS, which contains ethanol (resemblance of methanol), an alcohol known to permeabilize the membranes (Yuan, Xiong, Cohen, & Cohen, 2017) (Cell signaling Technology; https://www.cellsignal.com/learn-and-support/protocols/protocol-if-methanol, NovusBio; https://www.novusbio.com/support/fixation-and-permeabilization-in-icc-if).

Our work is not without limitations. First, when assessing the ease of manipulation of the slices, we did not quantify the number of slices that were torn and discarded. Second, although we targeted antigens present in different cells of the central nervous tissue, we did not test many other antigens that could be of interest in neuroscientific research, such as oligodendrocyte markers, CD markers, extracellular matrix markers, and other myelin antigens (e.g. myelin-associated glycoprotein or myelin oligodendrocyte glycoprotein) (Lyck et al., 2008). Also, we could have performed an immunofluorescence analysis to better depict antibody penetration through confocal microscopy. However, we chose IHC since it keeps well over time (storable), which is useful for further quantitative analysis, while immunofluorescence degrades with light exposure (photobleaching) (Im et al., 2019). IHC was also our staining of choice because it is routinely used by neuroscientists for antigen quantifications (e.g. demyelinating diseases) (Kuhlmann et al., 2017; Lassmann, 2018). Finally, we could also have assessed other basic colorations (e.g. luxol fast blue or cresyl violet), but we considered the assessment of the degradation of specific antigens as a more relevant target given their use in quantitative neuroscientific research and the impact of chemicals on antigen conformation (i.e. cross-linking) (Aoki, Kobayashi, & Isaki, 1999; Eltoum et al., 2001; Kuhlmann et al., 2017; Lyck et al., 2008; Puchtler & Meloan, 1985; Yamaguchi & Shen, 2013; Zhou et al., 2017).

To conclude, our work is the first to compare the quality and antigenicity of mice brain tissue fixed with solutions currently used in human gross anatomy laboratories. Even if FAS, routinely used in brain banks and animal fixation, produces better histology quality, we have successfully shown that antigenicity of four cell populations is preserved with two solutions used by anatomists. This opens the door to the use of human brains fixed with SSS and AS to assess different diseases using IHC. Further studies will test the same variables in human brain samples, optimizing the IHC protocol of SSS specimens to reduce the darkness of the background. These studies will also consider the post-mortem delay as a confounding variable, to evaluate the most suitable fixation method for both human dissection and research.

## Acknowledgements

We would like to thank Natural Sciences and Engineering Research Council of Canada grants to the JM and DB.

